# The contributions of the actin machinery to endocytic membrane bending and vesicle formation

**DOI:** 10.1101/172072

**Authors:** Andrea Picco, Wanda Kukulski, Hetty E. Manenschijn, Tanja Specht, John A. G. Briggs, Marko Kaksonen

**Affiliations:** Department of Biochemistry and NCCR Chemical Biology, University of Geneva, Science II, Quai Ernest-Ansermet 30, 1211 Geneva, Switzerland; Cell Biology and Biophysics Unit, European Molecular Biology Laboratory, Meyerhofstr.1, 69117 Heidelberg, Germany; Cell Biology Division, MRC Laboratory of Molecular Biology, Francis Crick Avenue, Cambridge CB2 0QH, United Kingdom; Structural and Computational Biology Unit, European Molecular Biology Laboratory, Meyerhofstr.1, 69117 Heidelberg, Germany; Structural Studies Division, MRC Laboratory of Molecular Biology, Francis Crick Avenue, Cambridge CB2 0QH, United Kingdom

**Author notes:** corresponding authors: Wanda Kukulski, Marko Kaksonen. equally contributing authors: Andrea Picco Wanda Kukulski.

## Abstract

Branched and crosslinked actin networks mediate cellular processes that move and shape membranes. To understand how actin contributes during the different stages of endocytic membrane reshaping, we analysed deletion mutants of yeast actin network components using a hybrid imaging approach that combines live imaging with correlative microscopy. We could thereby temporally dissect the effects of different actin network perturbations, revealing distinct stages of actin-based membrane reshaping. Our data show that initiation of membrane bending requires the actin network to be physically linked to the plasma membrane and to be optimally crosslinked. Once initiated, the membrane invagination process is driven by nucleation and polymerization of new actin filaments, independently of the degree of cross-linking and unaffected by a surplus of actin network components. A key transition occurs 2 seconds before scission when the filament nucleation rate drops. From that time point on, invagination growth and vesicle scission are driven by an expansion of the assembled actin network. The expansion is sensitive to the amount of filamentous actin and its crosslinking. Our results suggest that the mechanism by which actin reshapes the membrane adapts to force requirements that vary during the progress of endocytosis.

## Introduction

Arp2/3-mediated actin filament networks play key roles in generating and controlling force for movement and reshaping of cellular membranes (Rotty et al., 2013). These cytoskeletal assemblies drive diverse processes such as cell motility at the leading edge, formation of lamellipodia, generation of actin comet-tails for the intracellular movement of endosomes and pathogens (Taunton et al., 2000, Pollard and Borisy, 2003, Welch and Way, 2013), as well as several endocytic uptake pathways including clathrin-mediated endocytosis (Pelkmans et al., 2002, Galletta et al., 2010, Collins et al., 2011, Boulant et al., 2011). These fundamental cellular functions rely on force generation by treadmilling of actin monomers through a dendritic actin network built from a set of conserved protein components. The network architecture and dynamics are modulated through its composition (Svitkina, 2013). Thus, actin network properties can adjust to meet the requirements of different processes. However, it remains poorly understood how variations in network properties control the mechanisms of force generation in cells, and how the force requirements vary during actin-driven processes.

Endocytosis in budding and fission yeasts is strictly dependent on Arp2/3-based actin polymerization (Kubler and Riezman, 1993, Kaksonen et al., 2003, Galletta and Cooper, 2009), which provides force that drives membrane invagination against membrane tension and against the turgor pressure inside yeast cells (Aghamohammadzadeh and Ayscough, 2009, Basu et al., 2014). The process generally conforms to the dendritic nucleation model (Berro et al., 2010, Sirotkin et al., 2010). The membrane is coupled to actin through endocytic adaptors at the tip of the invagination, while nucleation and filament elongation occur at the plasma membrane (Kaksonen et al., 2003, Idrissi et al., 2012, Skruzny et al., 2012, Chen and Pollard, 2013, Berro and Pollard, 2014, Picco et al., 2015). Most molecular constituents of the yeast endocytic actin network have homologs in mammalian Arp2/3-mediated actin networks (Boettner et al., 2011a).

In this study, we aimed at mechanistically understanding how the endocytic actin network contributes throughout the process of membrane deformation and vesicle formation in the budding yeast *Saccharomyces cerevisiae*. We therefore deleted genes that encode proteins involved in coupling actin to the membrane, actin filament crosslinking, or regulation of actin nucleation. We studied these mutants using a hybrid imaging approach, which makes use of correlative microscopy data and of live cell imaging and particle tracking data (Kukulski et al., 2016, Picco et al., 2015, Picco and Kaksonen, 2017, Kukulski et al., 2011, Kukulski et al., 2012a). This approach provides measurements of membrane morphology and actin network volume as well as high-resolution information on protein localisation and assembly dynamics. We could thus observe and quantify very subtle phenotypes caused by specific actin network perturbations. Our previously published wild type data sets served us as reference data for interpreting the results presented here (Kukulski et al., 2012a, Picco et al., 2015). Our findings reveal three subsequent stages in the actin network lifetime suggesting that different actin based mechanisms may function during endocytic membrane shaping.

## Results

### Initiation of membrane bending requires actin-membrane coupling via clathrin adaptor proteins

The molecular mechanisms that form the initial bending of the endocytic membrane *in vivo* are unknown. While our previous data indicated that initiation of membrane invagination coincides with the assembly of actin, and inhibiting actin polymerization with Latrunculin A prevents membrane bending at endocytic sites (Kukulski et al., 2012a, Picco et al., 2015), immunoelectron microscopy data suggested that curvature of the endocytic membrane is induced before actin network assembly (Idrissi et al., 2012). Indeed, endocytic adaptor proteins could modulate membrane curvature before actin assembly through their membrane binding domains or through crowding, as demonstrated *in vitro* (Boucrot et al., 2012, Skruzny et al., 2015, Busch et al., 2015, Stachowiak et al., 2012). To shed light on the requirements for initiation of membrane bending we addressed the role of adaptor proteins under conditions of unhindered actin assembly. Epsins and Sla2 (the yeast homolog of Hip1R), act cooperatively to couple the membrane to the actin network via their membrane- and actin-binding domains, and are essential for endocytic vesicle formation (Kaksonen et al., 2003, Sun et al., 2005, Boettner et al., 2011b, Skruzny et al., 2012). We deleted the actin-binding domains of both Sla2 and Ent1 (*sla2ΔTHATCH ent1ΔACB-GFP*) to test if the presence of the membrane binding-domains of these endocytic adaptor proteins is sufficient for initiating membrane bending when the transmission of the force from the actin cytoskeleton is uncoupled from the plasma membrane (Skruzny et al., 2012). Cells carrying these mutations fail to form endocytic vesicles (Skruzny et al., 2012). By applying correlative light microscopy and electron tomography to these cells, we targeted 12 sites marked by the colocalisation of GFP-labeled epsin and Abp1-mCherry as a marker for the presence of an actin network. In all the target localisations, we found no invaginations but only flat plasma membranes associated with large cytoplasmic zones excluding ribosomes (Figure 1A). In a similar experiment on cells in which the whole *SLA2* gene was deleted, all 11 target localisations of Sla1-GFP and Abp1-mCherry showed also flat plasma membranes associated with large exclusion zones (Figure 1B, and Table 1). The ribosome exclusion zones represent the extent of the endocytic machinery including the actin network (Kukulski et al., 2012a). In wild type cells, the presence of the actin network always coincides with membrane invaginations or vesicles (Kukulski et al., 2012a). Therefore, these results indicate that actin polymerization and the presence of all membrane-binding domains are not sufficient to induce membrane invagination, but the two need to be physically linked (Figure 1C). Coupling the membrane to a polymerizing actin network via the actin-binding domains of Sla2 and Ent1 is thus essential for initiating endocytic membrane invagination.

**Figure 1:**
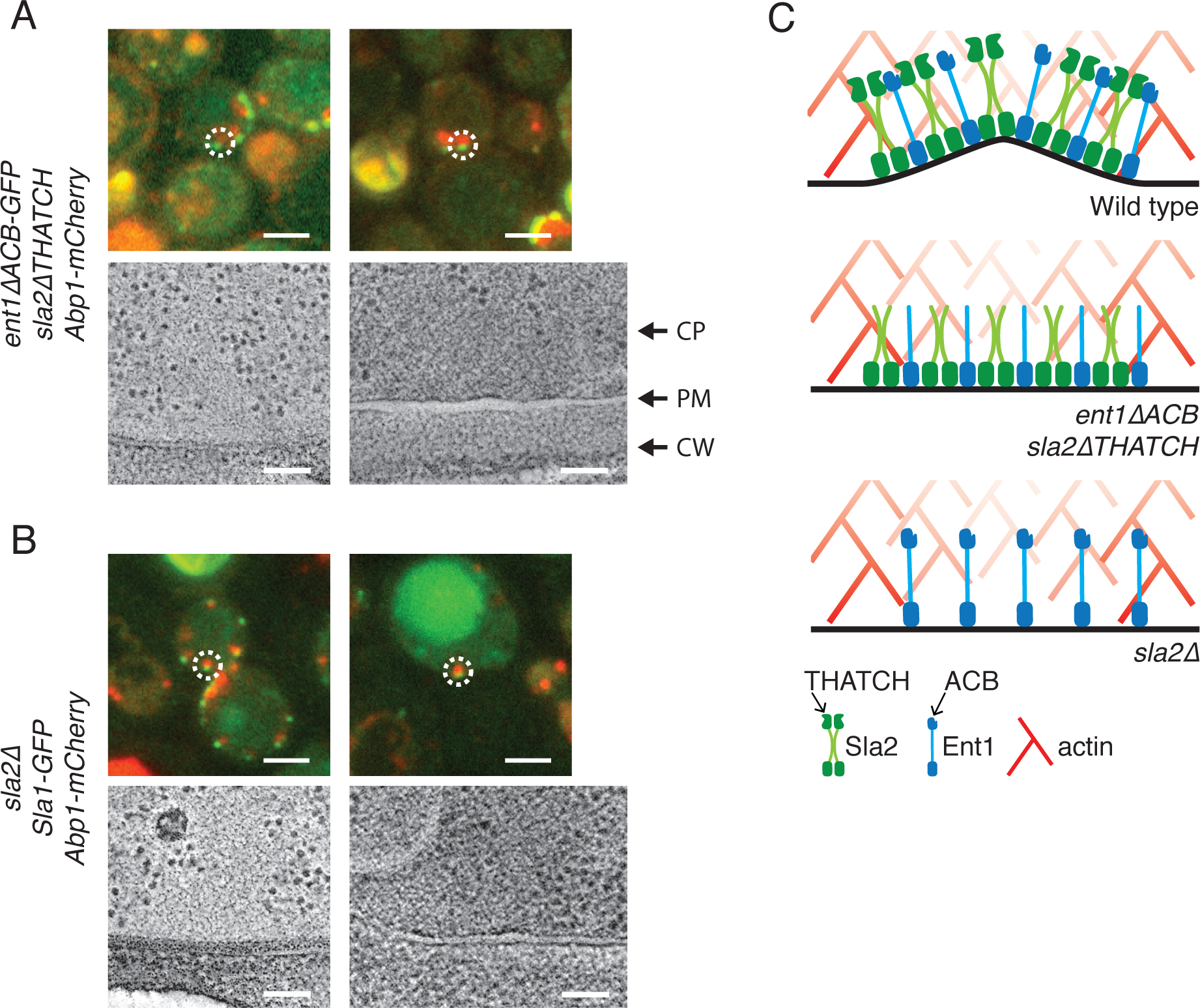
Correlative microscopy of endocytosis in cells with the actin cytoskeleton uncoupled from the plasma membrane. (**A**) Upper row shows overlaid red and green channel fluorescence images of resin sections of yeast cells expressing *ent1ΔACB-GFP*, *sla2ΔTHATCH* and Abp1-mCherry. Endocytic sites targeted by correlative microscopy are marked by white dashed circles. Lower row shows virtual slices from electron tomograms at the corresponding localisations. (**B**) Upper row shows overlaid red and green channel fluorescence images of resin sections of *sla2Δ* yeast cells expressing Sla1-GFP and Abp1-mCherry. Endocytic sites targeted by correlative microscopy are marked by white dashed circles. Lower row shows virtual slices from electron tomograms at the corresponding locations. In both A and B, the panels to the left show examples of the flattest membranes in each dataset. The panels on the right show examples of the most bent membranes in each dataset. The dataset sizes were n = 12 for (A), n= 11 for (B). All panels are oriented so that the cytoplasm (CP) is above the plasma membrane (PM), and the cell wall (CW) is below. Scale bars are 2 µm in fluorescence images, 100 nm in electron tomography images. (**C**) Model representation for the assembly of Sla2, Ent1 and the actin cytoskeleton. In wild type cells, assembly of intact Sla2, Ent1 and actin leads to bending of the membrane. In absence of the actin binding domains of Sla2 and Ent1, THATCH and ACB, respectively, or in the absence of full-length Sla2, the membrane remains flat despite actin polymerization.

**Table 1:** Sample sizes of correlative microscopy data.

| Yeast strain | Data set size: number of endocytic structures obtained from CLEM data | Number of individual fluorescent signals analysed by CLEM | number of tomograms acquired to obtain the data set | number of EM grids imaged by CLEM to obtain the data set | number of HPF samples (i.e. yeast pellets in resin block) used to obtain the data set |
| --- | --- | --- | --- | --- | --- |
| sla2delta, Sla1-EGFP, Abp1-mCherry | 11 | 11 | 6 | 2 | 1 |
| ent1deltaACB-GFP, sla2deltaTHATCH, Abp1-mCherry | 12 | 12 | 9 | 2 | 1 |
| sac6delta, Sla1-EGFP, Abp1-mCherry | 60 | 59 | 41 | 6 | 1 |
| bbc1delta, Sla1-EGFP, Abp1-mCherry | 58 | 52 | 35 | 3 | 2* |
| bbc1delta, Rvs167-EGFP, Abp1-mCherry | 37 | 29 | 21 | 2 | 1 |

\*from two different cell cultures

### Actin filament crosslinking is critical for the initiation of membrane bending and for reaching scission stage

The composition of actin networks modulates their mechanical properties. For instance, the stiffness of actin gels decreases with a lower density of actin filament crosslinkers (Blanchoin et al., 2014). In yeasts, the actin crosslinking activity of fimbrin has been shown to be essential for endocytic internalization (Kubler and Riezman, 1993, Kaksonen et al., 2005, Skau et al., 2011). We perturbed actin filament crosslinking by deleting the yeast fimbrin Sac6 and studied *sac6*Δ cells by live imaging and correlative microscopy to investigate the effects on membrane invagination and vesicle formation.

We first tracked the dynamic behaviour of the coat protein Sla1-GFP in *sac6Δ* cells with high spatiotemporal resolution (Picco and Kaksonen, 2017). Sla1 is positioned close to the invagination tip and can be used to track the movement of the membrane invagination and the vesicle into the cell (Kukulski et al., 2016, Picco et al., 2015, Idrissi et al., 2008). In *sac6Δ* cells, we found three distinct types of Sla1-GFP behaviour (Figure 2A). The majority of Sla1 spots (78.4% ± 5.5%, mean ± SE, 145 endocytic events in 9 cells; Figure 2A; see ‘Materials and methods’ and Table 2) remained immobile at the plasma membrane until they disassembled, consistent with earlier studies (Kaksonen et al., 2005, Gheorghe et al., 2008). However, 15.1% ± 4.4% (mean ± SE, 145 endocytic events in n = 9 cells) of events revealed a distinct behaviour: The initial phase of centroid movement was very similar to that in wild type cells,but at a position approximately 90 nm away from the plasma membrane, the centroid started to retract towards the plasma membrane (Figure 2A and B). The remaining 6.5% ± 2.2% (mean ± SE, 145 endocytic events in n = 9 cells) of Sla1-GFP spots were motile and moved inward similarly to the Sla1-GFP wild type cells (Figure 2A and C).

**Figure 2:**
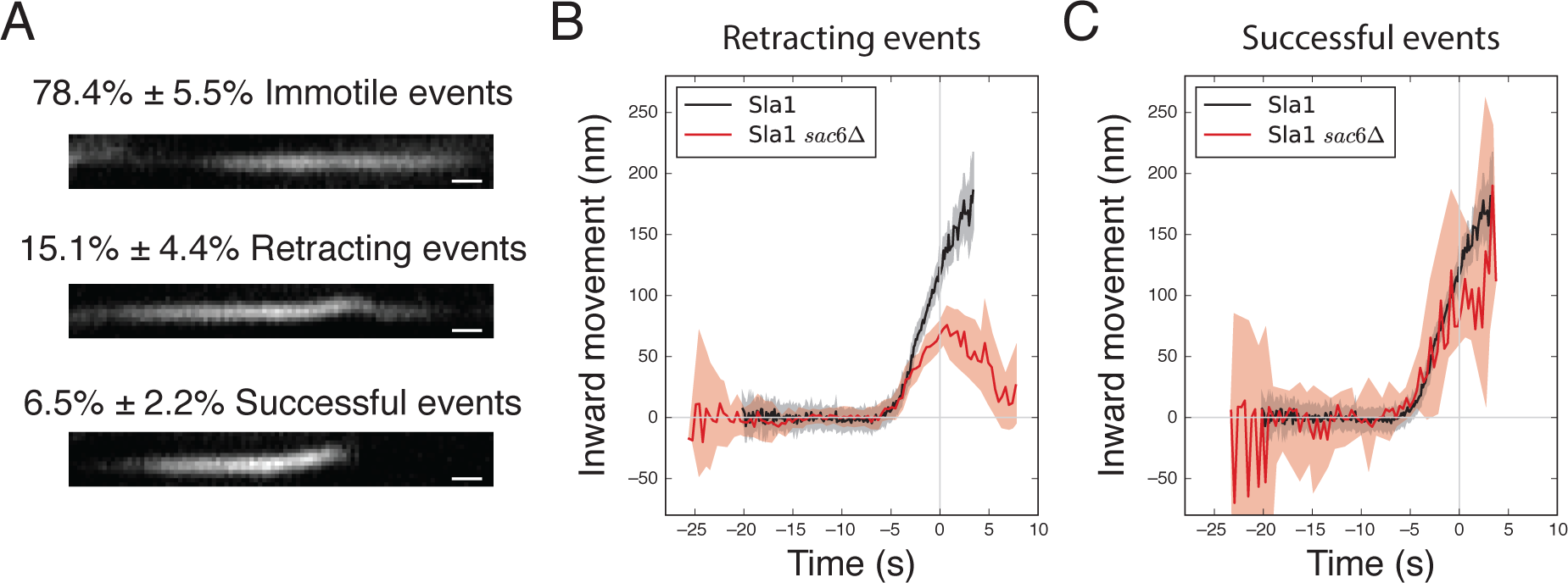
Live imaging of endocytosis in cells with impaired actin crosslinking. (**A**) Representative kymographs of the three distinct behaviours of Sla1-GFP spots in *sac6Δ* cells and the percentage of occurrence for each type of event (mean ± SE, n = 9 cells). Scale bars correspond to 5 seconds. (**B**) The inward movement of Sla1-GFP average trajectories of retracting events in *sac6Δ* cells (red) and of wild type cells (black, (Picco et al., 2015)). (**C**) The inward movement of Sla1-GFP average trajectories of successful events in *sac6Δ* cells (red) and of wild type cells (black, (Picco et al., 2015)). The shadows in (**B**) and (**C**) correspond to the 95% confidence interval. Time 0 in (**B**) and (**C**) marks the scission event (Kukulski et al., 2012a, Picco et al., 2015).

To visualize the membrane morphology directly, we applied correlative microscopy to *sac6Δ* cells expressing Sla1-EGFP and Abp1-mCherry. In wild type cells, the presence of these proteins marks events that span from initiation of membrane bending until disassembly of the actin network from the newly formed endocytic vesicle (Kukulski et al., 2012a). In *sac6Δ*, we found that, in the presence of both Abp1 and Sla1, as well as in the presence of Abp1 only, the majority of endocytic sites were flat membranes (72% and 62%, n = 25 and n =34, respectively) and the remaining ones were invaginations (28% and 38%, n = 25 and n =34, respectively) (Figure 3A and B). We did not find endocytic vesicles at any of the sites, confirming that successful scission events are very rare in *sac6Δ* cells. These data also support the live cell imaging observation that the majority of the Sla1-GFP spots remained immobile throughout their lifetime. The immobile spots are thus likely to be events in which membrane bending is not initiated, despite assembly of the actin network.

**Figure 3:**
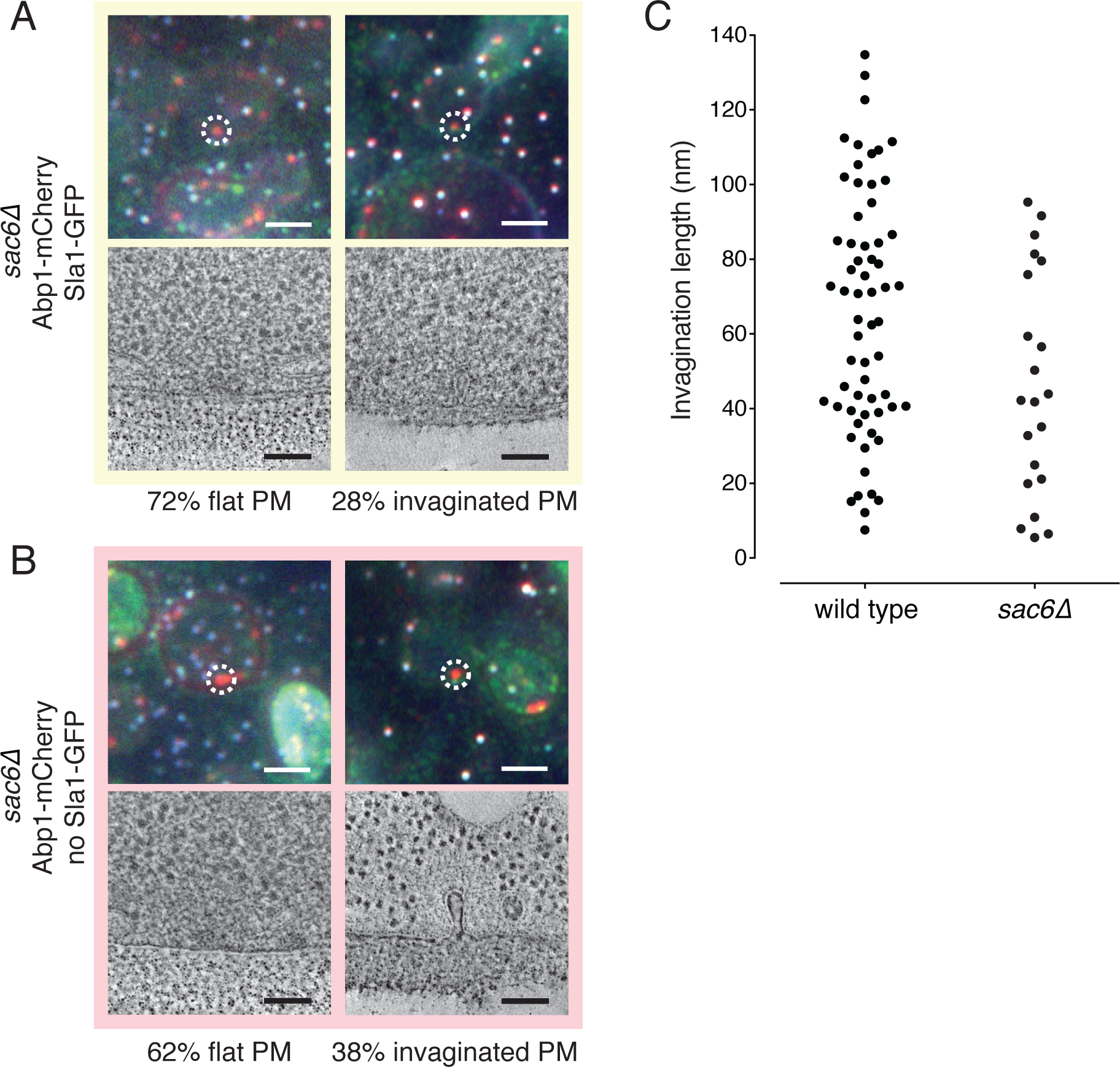
Correlative microscopy of endocytosis in cells with impaired actin crosslinking. (**A**) Upper row shows overlaid red, green and blue channel fluorescence images of resin sections through *sac6Δ* yeast cells expressing Abp1-mCherry and Sla1-GFP. The blue channel indicates TetraSpeck fiducial markers for correlation. White dashed circles mark endocytic sites where Abp1-mCherry and Sla1-GFP colocalize, targeted by correlative microscopy. Lower row are virtual slices from electron tomograms at the corresponding localisations. Of n = 25 sites, 72% displayed flat plasma membranes, and 28% showed invaginations. (**B**) Upper row shows overlaid red green and blue channel fluorescence images of resin sections through yeast cells. The blue channel indicates TetraSpeck fiducial markers for correlation. White dashed circles mark endocytic sites where Abp1-mCherry is present but Sla1-GFP is absent, targeted by correlative microscopy. Lower row shows virtual slices from electron tomograms at the corresponding localisations. Of n = 34 sites, 62% displayed flat plasma membranes, and 38% showed invaginations. Scale bars are 2 µm in fluorescence images, 100 nm in electron tomography images. (**C**) The distribution of endocytic invagination lengths measured in the electron tomograms of *sac6Δ* cells. The wild type data (Kukulski et al., 2012a) is shown for comparison.

**Table 2:** Sample sizes of live imaging data.

|  | Protein | Number of single color trajectories used to generate the average trajectory | Number of cells imaged to obtain the single color trajectories | Number of trajectory pairs used for spatial and temporal alignment | Number of cells imaged to obtain the trajectory pairs |
| --- | --- | --- | --- | --- | --- |
| bbc1delta | Abp1 | 52 | 13 | NA | NA |
| bbc1delta | Rvs167 | 65 | 14 | 56 | 21 |
| bbc1delta | Sla1 | 49 | 13 | 47 | 13 |
| bbc1delta | Arc18 | 51 | 15 | 61 | 20 |
| bbc1delta | Las17 | NA | NA | 50 | 19 |
| sac6delta | Sla1 motile | 7 | 15 | NA | NA |
| sac6delta | Sla1 retracting | 19 | 15 | NA | NA |
| Wild type | Abp1 | 66 | 13 | NA | NA |
| Wild type | Rvs167 | 60 | 13 | 262 | 69 |
| Wild type | Sla1 | 67 | 13 | 58 | 14 |
| Wild type | Arc18 | 70 | 14 | 90 | 12 |
| Wild type | Las17 | NA | NA | 49 | 15 |

We then measured the lengths of the invaginations marked by presence of Abp1-mCherry and Sla1-GFP, or by presence of Abp1-mCherry and absence of Sla1-GFP, in *sac6Δ* cells. We found that all the invaginations were less than 100 nm long, while in wild type cells invaginations are up to 140 nm long (Kukulski et al., 2012a) (Figure 3C). These measurements are in agreement with the live imaging finding that most invaginations retract before reaching 100 nm in length (Figure 2B).

We also measured the curvature of invagination tips in *sac6Δ* cells. Like in wild type cells, the tips reached a minimum radius for invaginations of about 40 nm in length. However, the average tip radius for invaginations longer than 40 nm was significantly larger in *sac6Δ* cells than in wild type cells (*sac6*Δ: 15.5 nm, SD 3.7 nm, n=12, wt: 11.6 nm, SD 2.6 nm, n=47; p=0.0003) (Supplemental Figure S1). Thus, wider tips are either a property of all invaginations in *sac6Δ* cells, or widening of invagination tips occurs during retraction, possibly due to disassembly of coat proteins.

Taken together, our results indicate that in absence of the actin crosslinker Sac6, endocytosis is blocked either at the stage of initial membrane bending, or at an invagination length of approximately 100 nm, followed by a retraction of the invagination. Remarkably, in those events that initiate membrane bending, elongation of the invagination appears to be unaffected until retraction occurs. In rare cases, endocytosis progresses further and likely leads to successful vesicle formation.

### Perturbed actin nucleation in *bbc1*Δ cells enhances propulsion of vesicles

Arp2/3-dependent nucleation of actin filaments is driven by nucleation promoting factors (NPFs) at cellular membranes (Pollard, 2007, Campellone and Welch, 2010). The NPF Las17, a yeast N-WASP ortholog, is required for actin assembly at endocytic sites and its activity is negatively regulated by Bbc1 *in vitro* (Rodal et al., 2003, Sun et al., 2006). At endocytic sites, Bbc1 colocalises with actin spots (Kaksonen et al., 2005). We perturbed the rate of actin filament nucleation by deleting *BBC1*. In *bbc1*Δ cells, endocytic actin spots are brighter and the endocytic coat moves inward faster and over a longer distance than in wild type cells (Kaksonen et al., 2005).

To understand how actin nucleation in the absence of Bbc1 affects invagination dynamics, we first tracked Sla1-GFP dynamics in *bbc1Δ* cells. The average centroid trajectory of Sla1-GFP in *bbc1Δ* cells showed an initial movement at the same speed as in wild type cells. About 7 seconds later, approximately when scission occurs in wild type cells, Sla1 centroid movement accelerated and it moved deeper into the cell than in the wild type (Figure 4A). We next investigated the dynamics of the amphiphysin Rvs167 in *bbc1Δ* cells. In wild type cells, the peak of Rvs167-GFP fluorescence intensity marks the scission event and coincides with a fast inward movement of the Rvs167-GFP centroid (Picco et al., 2015). In *bbc1*Δ cells, Rvs167-GFP showed a pronounced movement into the cytoplasm, which was concomitant with its peak in number of molecules (Figure 4B and Supplemental Figure S2A), in 85% ± 6% (median ± SE, 195 endocytic events in n = 14 cells) of the endocytic events. Like Sla1-GFP, motile Rvs167GFP spots also moved deeper into the cell cytoplasm (Figure 4B). In the remaining 15% ± 6% (median ± SE, 195 endocytic events in n = 14 cells) of events, Rvs167-GFP spots appeared immotile. The trajectories of Sla1-GFP and Rvs167-GFP indicate that *bbc1Δ* does not significantly affect the rate of invagination growth. However, they suggest that, after scission, vesicles are moved deeper into *bbc1Δ* cells compared to wild type.

**Figure 4:**
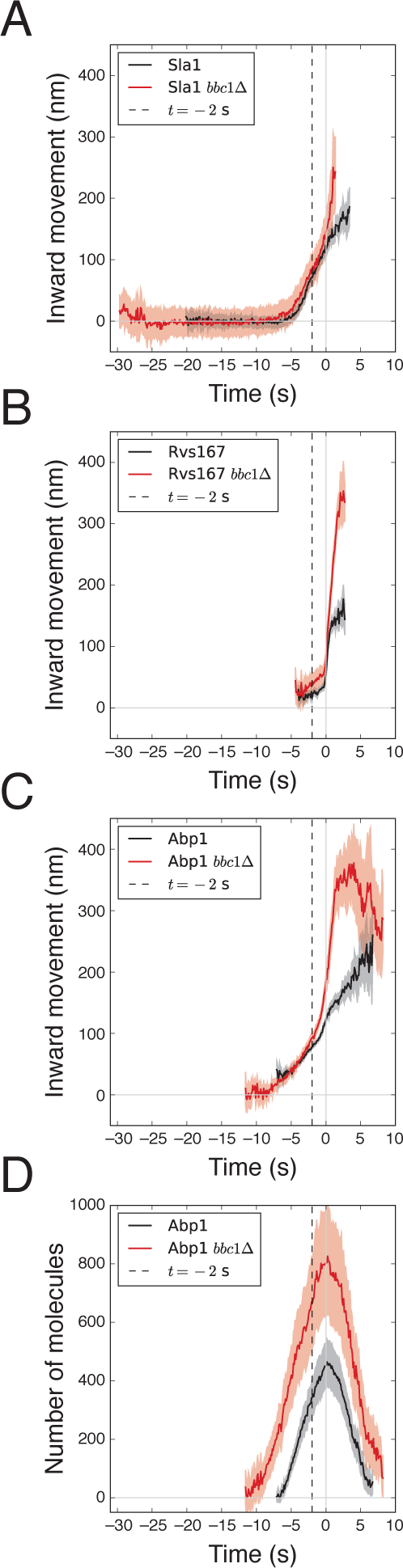
Live imaging of endocytosis in cells with perturbed actin nucleation. (**A**) The inward movement of Sla1-GFP average trajectories in *bbc1*Δ (red) and in wild type (black) cells.(**B**) The inward movement of Rvs167-GFP average trajectories in *bbc1Δ* (red) and wild type (black) cells. (**C**) The inward movement of Abp1-GFP average trajectory in *bbc1Δ* (red) and in wild type (black) cells. (**D**) The number of Abp1-GFP molecules over time in *bbc1Δ* (red) and in wild type (black) cells. The shadows in A-D correspond to the 95% confidence interval. The trajectories in *bbc1Δ* and wild type cells were independently aligned in space and in time to Abp1 (Picco and Kaksonen, 2017). The 0 on the Y-axis marks the position of Sla1-GFP prior the invagination movement starts. The 0 on the X-axis marks the scission event (see Materials and Methods, Image analysis and trajectory alignment; (Kukulski et al., 2012a, Picco et al., 2015, Picco and Kaksonen, 2017)). The wild type trajectories are from (Picco et al., 2015).

We then imaged Abp1-GFP, which we used as a marker for the dynamics of the endocytic actin network (Kaksonen et al., 2003). In wild type cells, the recruitment of Abp1 coincides with the start of the Sla1 inward movement, which marks membrane invagination (Picco et al., 2015). In *bbc1Δ* cells, the recruitment of Abp1 preceded the start of Sla1-GFP inward movement by about 4 seconds, indicating that actin assembly does not immediately lead to membrane bending. The centroid of Abp1-GFP initially localised closer to the plasma membrane than in wild type cells (Figure 4C), suggesting a different distribution of the actin filaments on the membrane. Interestingly, the Sla1 inward movement began when the actin network reached the same height as in wild type cells.

The endocytic actin spots are brighter in *bbc1Δ* cells (Kaksonen et al., 2005). We therefore measured the number of Abp1-GFP and Arc18-GFP molecules present at endocytic sites over time (Joglekar et al., 2006, Lawrimore et al., 2011, Picco et al., 2015) (Figure 4D and Supplemental Figure S2B, see Materials and Methods). Abp1 and Arc18 were recruited to the endocytic sites at the same rate as in wild type cells (Picco et al., 2015). However, because assembly of the actin network in *bbc1Δ* cells lasted for about 5 seconds longer than in wild type cells, the fully-assembled network contained more Abp1 and Arc18 molecules. At the time when scission occurred, the number of Abp1 molecules recruited to the endocytic site was about doubled (Figure 4D), while the number of Arc18 was about 25% larger compared to wild type (Supplemental Figure S2B).

The Abp1-GFP centroid in *bbc1Δ* cells began moving inward at the same speed as in wild type cells. However, the speed of the Abp1 centroid drastically increased when the invagination length reached 100 nm, about 2 seconds before scission (Figure 4C). Interestingly, the speed-up of the Abp1 centroid was not linked to an increase in assembly rate of Abp1 molecules (Figure 4D). This observation suggests that 2 seconds before scission, the actin network is undergoing a rearrangement, which leads to increased motility without extra addition of protein components.

We next performed correlative microscopy of *bbc1Δ* cells expressing Sla1-GFP and Abp1mCherry, as well as *bbc1Δ* cells expressing Rvs167-GFP and Abp1-mCherry. Similarly to wild type cells, we found endocytic invaginations and vesicles in the presence of Abp1 and Sla1 or Rvs167, as well as in the presence of Abp1 but the absence of Sla1 or Rvs167 (Figure 5A and B). We measured that endocytic vesicles were positioned up to 600 nm from the plasma membrane, which is significantly further away from the plasma membrane than in wild type cells (*bbc1Δ*: 288.7 nm, SD: 158.7 nm, n=47; wt: 129.5 nm, SD: 39.7 nm, n=62; P<0.0001) (Figure 5C). This result is in striking agreement with the enhanced movement of Sla1-GFP and Rvs167-GFP starting around scission. Taken together, these results suggest that vesicles are being propelled faster and deeper into the cytoplasm upon deletion of *bbc1Δ*.

**Figure 5:**
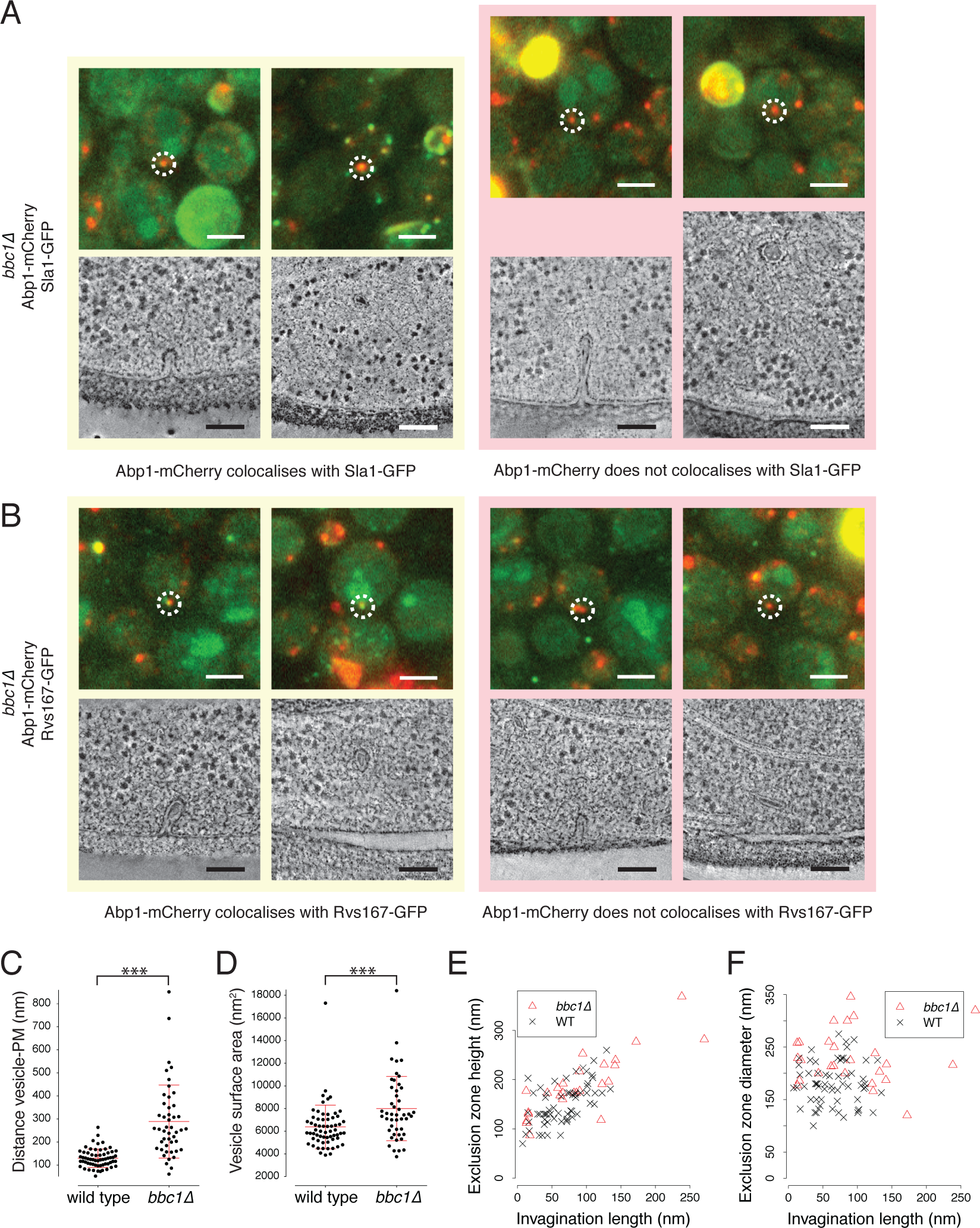
Correlative microscopy of endocytosis in cells with perturbed actin nucleation. (**A**) *bbc1Δ* cells expressing Abp1-mCherry and Sla1-GFP. Upper row shows overlaid red and green channel fluorescence images of resin sections through yeast cells. White dashed circles mark endocytic sites where Abp1-mCherry and Sla1-GFP colocalize (yellow box), or where Abp1mCherry is present but Sla1-GFP is absent (pink box), targeted by correlative microscopy. Lower row are virtual slices from electron tomograms at the corresponding localisations. (**B**) *bbc1Δ* cells expressing Abp1-mCherry and Rvs167-GFP. Upper row shows overlaid red and green channel fluorescence images of resin sections through yeast cells. White dashed circles mark endocytic sites where Abp1-mCherry and Rvs167-GFP colocalize (yellow box), or where Abp1-mCherry is present but Rvs167-GFP is absent (pink box), targeted by correlative microscopy. Lower row are virtual slices from electron tomograms at the corresponding localisations. Scale bars are 2 µm in fluorescence images, 100 nm in electron tomography images. (**C**) The shortest distance between vesicle centres and the plasma membrane (PM) in wild type and *bbc1Δ* cells. (**D**) The surface areas of vesicles in *bbc1Δ* cells. Red lines in C and D represent the mean and the standard deviation. (**E**) The heights of exclusion zones surrounding invaginations plotted against invagination lengths. (**F**) The diameters of exclusion zones surrounding invaginations plotted against invagination lengths. Data from wild type cells in C F are from (Kukulski et al., 2012a).

To better resolve the effects of *bbc1Δ* on vesicle formation, we measured the surface areas of endocytic vesicles and compared them to wild type. We found that vesicles in *bbc1*Δ cells were on average 25% larger (*bbc1Δ*: 7’998 nm^2^, SD: 2’838 nm^2^, n=47, wt: 6’380 nm^2^, SD: 1’929 nm^2^, n=62; P=0.0004) (Figure 5D). The larger vesicle sizes indicate that scission either occurred when invaginations were longer, or that scission sites on the invaginations were positioned closer to the plasma membrane than in wild type cells. The putative sites of scission are the necks of invaginations, which we found to be positioned similarly to those in wild type cells (Supplemental Figure S2C). However, we found that there were longer invaginations in *bbc1Δ* cells compared to the wild type (Supplemental Figure S2D), suggesting that the membrane inward movement, observed through centroid tracking of Sla1-GFP (Figure 4A), might accelerate already shortly before scission, leading to longer invaginations that result in larger vesicles.

### Actin assembly occurs on a larger membrane area in *bbc1*Δ cells

To detail whether the increase in Abp1 molecules is accompanied by an increase in actin network size in *bbc1Δ* cells, we measured the dimensions of the ribosome exclusion zones surrounding invaginations. We found that the volumes of exclusion zones in *bbc1Δ* cells were significantly larger compared to those in wild type cells (*bbc1Δ*: 0.0052 fl, SD=0.0029 fl, n=21. wt: 0.0028 fl, SD=0.0017 fl, n=62; P=0.0001), which was in line with the increased amounts of Abp1-GFP molecules that we measured by fluorescence microscopy. Interestingly, while the heights of the exclusion zones in *bbc1Δ* correlated with the invagination lengths as observed in wild type cells (Figure 5E) (Kukulski et al., 2012a), the exclusion zone diameters were independent of the stage of the invagination process and significantly larger than in wild type cells (Figure 5F, *bbc1Δ*: mean=235.1 nm, SD=48.2 nm, n=21, wt: mean=181.2 nm, SD=40.9 nm, n=62; P<0.0001). Note that to compute the above values we only considered exclusion zones of *bbc1Δ* invaginations below 135 nm in length so that the invagination lengths compare with the wild type population. The enlarged diameters of the exclusion zones indicate that the actin cytoskeleton is distributed over a larger surface area of the plasma membrane when nucleation of actin is initiated. These results are consistent with our observation that the initial position of the Abp1-GFP centroid in *bbc1Δ* cells is closer to the plasma membrane than in wild type cells (Figure 4C). Therefore, despite the height of the actin network being comparable to wild type during the invagination process, the volume of the actin network is larger.

Like in wild type cells, vesicles in *bbc1Δ* cells were surrounded by ribosome exclusion zones that were either disconnected or connected to the plasma membrane (Figure 5A and B). We previously suggested that the latter represent the earliest phase of the vesicle lifetime (Kukulski et al., 2012a). In *bbc1Δ* cells, only 20% of vesicles had exclusion zones connected to the plasma membrane, as compared to 70% in wild type cells. Therefore, we conclude that the density of the actin network near the plasma membrane decreases earlier in *bbc1Δ* cells than in wild type cells. This asymmetric change in density could shift the centroid of the Abp1GFP towards the cell centre, contributing to enhance the movement of the Abp1-GFP centroid in *bbc1Δ* cells.

### Actin nucleation ceases before scission, while network expansion and movement continue in *bbc1*Δ cells

We next investigated the number of molecules of Las17 at the endocytic site. Las17 is a NPF and putative target of Bbc1 (Kaksonen et al., 2005, Rodal et al., 2003). Las17 accumulates on the plasma membrane, where it activates the Arp2/3 complex to nucleate actin filament formation (Sun et al., 2006, Winter et al., 1999). We found that in *bbc1Δ* cells, Las17 accumulated over a longer time than in wild type cells (Figure 6). In wild type cells, the number of Las17 molecules plateaus for about 5 seconds at about 40 molecules and then the number of molecules starts decreasing about 2 seconds before scission (Picco et al., 2015). In *bbc1Δ*, however, the number of Las17 molecules did not plateau but continued accumulating until it reached about 80 molecules, and then started to disassemble about 2 seconds before scission like in wild type cells (Figure 6). The assembly of the actin cytoskeleton in *bbc1Δ* cells began when approximately 40 Las17 molecules had accumulated, like in wild type cells. Taken together, these data show that in *bbc1Δ* cells, Las17 accumulation is unhindered and the number of accumulating Las17 molecules is larger than in wild type, supporting a role for Bbc1 in regulating the assembly of Las17. Further, the data suggest that Bbc1 is not required for triggering disassembly of Las17 and thereby for ending the accumulation of actin network components.

**Figure 6:**
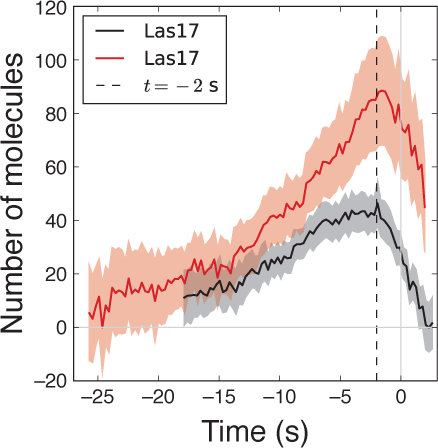
Live imaging of nucleation promoting factor in cells with perturbed actin nucleation. The number of Las17-GFP molecules over time in *bbc1Δ* (red) and in wild type (black) cells. The shadows correspond to the 95% confidence interval. Time 0 marks the scission event, which is timed according to the peak in the Rvs167 number of molecules, when aligned in time to Abp1 (see Materials and Methods, Image analysis and trajectory alignment; (Kukulski et al., 2012a, Picco et al., 2015)). The wild type data are from (Picco et al., 2015).

In summary, in *bbc1*Δ cells, the actin network starts accumulating on a larger membrane area than in wild type cells, resulting in larger network volumes. Nevertheless, the rate at which the actin network expands towards the cytoplasm is similar to wild type cells up to about 2 seconds before scission: The velocity of the actin centroid movement as well as the height of the exclusion zones are similar to wild type cells. In the last 2 seconds before scission occurs, the rate of actin nucleation drops and a rearrangement of the actin network results in a significant speed-up of the actin centroid movement, as compared to its initial movement and to its movement in wild type cells. Thus, the rate and progression of the invagination process are unaffected until shortly before scission. The changes in actin network behaviour seem to interfere only with the shape of late invaginations, which can get longer than in wild type cells and therefore result in larger vesicles, and with the movement of vesicles, which are pushed further into the cytoplasm.

## Discussion

The endocytic actin network is critical for invaginating the plasma membrane during endocytosis (Merrifield et al., 2002, Kaksonen et al., 2003, Boulant et al., 2011, Kukulski et al., 2012a). Here, we genetically manipulated different biochemical activities related to assembly and connectivity of the actin network to gain understanding of the molecular mechanisms underlying endocytic membrane bending.

Based on the mutant phenotypes that we studied here, we can define three sequential stages in the actin network lifetime (Figure 7). The first stage corresponds to assembly of the actin network until initiation of membrane bending. The second stage spans the time of invagination growth up to about 100 nm, 2 seconds before vesicle scission. During the second stage, the numbers of Arp2/3 and actin molecules increase, suggesting that the growth of the invagination is driven by nucleation and polymerization of actin filaments (Picco et al., 2015, Sirotkin et al., 2010, Berro et al., 2010). The third stage spans the last 2 s of the invagination and the formation of the endocytic vesicle. During the third stage, the numbers of Arp2/3 and actin molecules have plateaued and the numbers of the main NPFs, Las17 and Myo5, decrease, suggesting that the rate of filament nucleation drops (Picco et al., 2015). However, during the third stage the centroids of the actin cytoskeletal proteins keep moving inward, and the actin network volume increases, indicating that the network expands although the amount of actin is not increasing. These observations could mean that different mechanisms of propulsion are used during the second and third stages.

**Figure 7:**
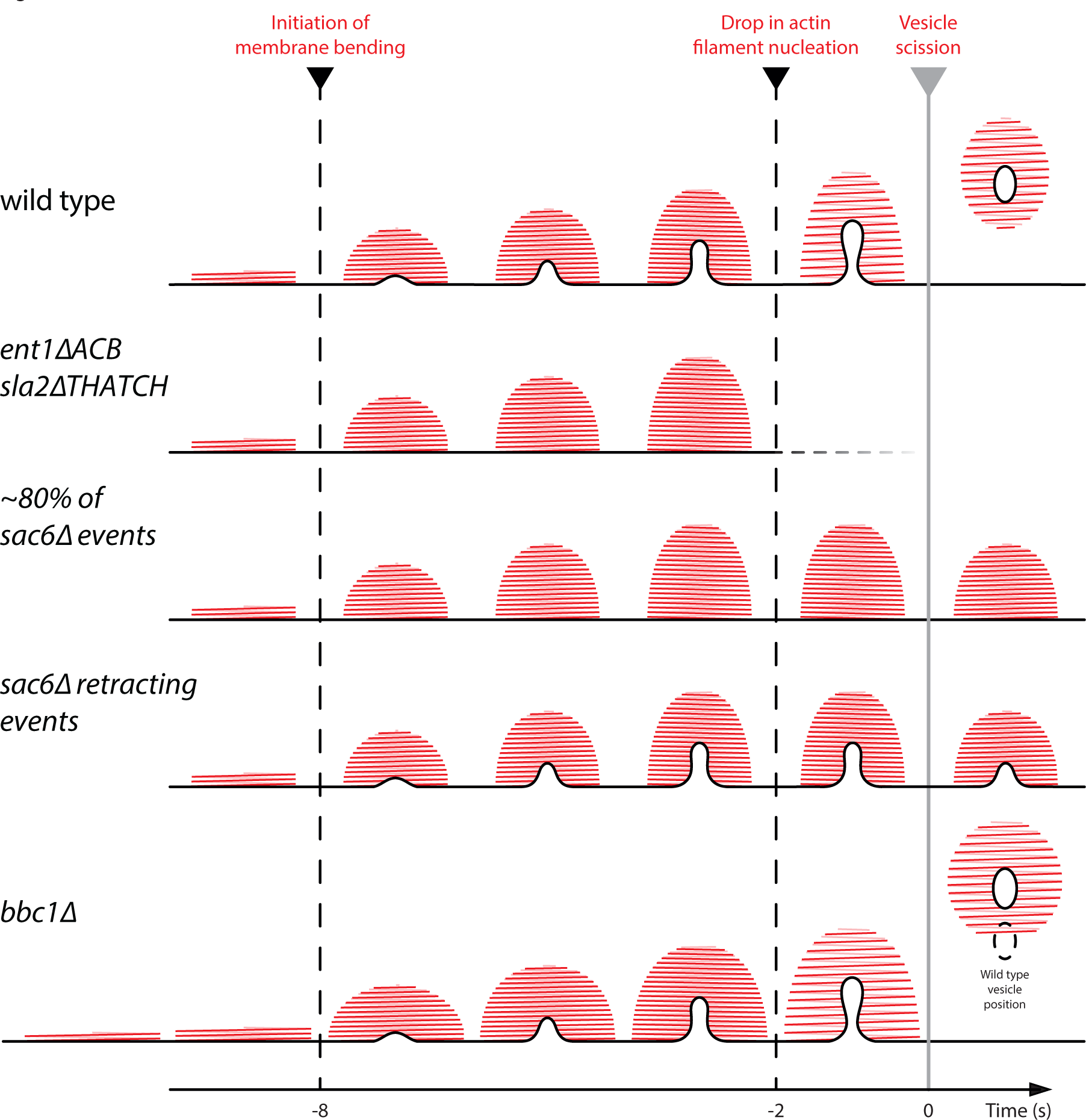
The actin network pulls the plasma membrane invagination in three distinct stages. The three stages correspond to 1) actin network assembly until the initiation of membrane bending is achieved, 2) invagination growth driven by actin nucleation and polymerization, and 3) expansion of the actin network after the nucleation of new actin filaments has dropped 2 seconds before scission. The transitions between the stages are indicated by the dashed lines. *ent1ΔACB*, *sla2ΔTHATCH* prevents initiation of membrane bending. Likewise, *sac6Δ* hinders initiation of membrane bending in the majority of cases. If membrane bending is initiated, the plasma membrane invagination proceeds until 2 seconds before scission and then retracts. In rare cases, invagination growth proceeds further through scission of the vesicle, like in wild type cells. *bbc1Δ* allows the plasma membrane to invaginate normally but the actin network is larger and expands about 2 seconds before scission. In *bbc1Δ* the vesicle is propelled deeper into the cytoplasm than in wild type cells.

Endocytosis is critically dependent on the actin-binding regions of the clathrin-adaptors Sla2 and Ent1 (Skruzny et al., 2012). Here we show that these actin-binding regions are critical for initiating membrane bending. If these regions are deleted the plasma membrane at the endocytic site remains flat (Figure 7). This result suggests that the assembly of the clathrin coat and the network of actin filaments are not sufficient to initiate membrane bending *in vivo* in yeast cells. The initiation of membrane bending requires that the plasma membrane is physically linked to the actin network via the clathrin adaptors.

In *sac6*Δ cells, most of the endocytic events fail to initiate the invagination process. Sac6mediated crosslinking of actin filaments is thus important for initial bending of the plasma membrane. In about a fifth of the events, membrane bending is initiated and the invaginations grow normally up to about 100 nm in length, which in wild type cells corresponds to about 2 seconds before scission (Picco et al., 2015). The second stage is thus unaffected in *sac6*Δ cells, if membrane bending is initiated. However, the majority of invaginations in *sac6*∆ cells retract and only very few events proceed through apparently normal scission of the vesicle. Therefore, crosslinking is likely to also be important for the expansion of the actin network, which characterizes the third stage that drives invagination growth through scission. In the rare successful events in *sac6Δ* cells, stochastic variations in the process, for example, the amount of other crosslinkers such as Scp1 (Aghamohammadzadeh and Ayscough, 2009), may allow the invagination to proceed through scission despite the absence of Sac6.

In *bbc1*Δ cells, more actin is assembled, which results in a wider actin network than in wild type cells. The start of membrane bending in *bbc1*Δ cells is delayed with respect to the initiation of actin assembly, suggesting that during the first stage, the actin network is not organized in an optimal way to produce force for invagination growth. However, as soon as membrane bending is initiated, the invaginations in *bbc1*∆ cells grow at the same rate as in wild type cells. During the third stage in *bbc1*∆ cells, the centroid movement of both actin cytoskeletal proteins and coat proteins is enhanced. This suggests that the final expansion or rearrangement of the actin network is strongly enhanced in *bbc1*∆ cells, and thereby accelerates the coat movement.

The force that is needed to pull a membrane tube depends on membrane shape and degree of protein coating (Powers et al., 2002, Koster et al., 2005, Derenyi et al., 2002, Dmitrieff and Nedelec, 2015). For endocytosis, it has been suggested that the force required for the initiation of membrane bending is higher than the force needed to subsequently shape the membrane into a tubular invagination (Dmitrieff and Nedelec, 2015). Our observations suggest that the formation of the actin network might have evolved to allow force generation to adjust to the changing force requirements during the different stages of endocytosis. The crosslinking of actin filaments controls the mechanical properties of the actin network: A network of crosslinked actin filaments sustains larger stresses without deformation and has a higher elastic modulus than actin filaments alone (Sato et al., 1987, Wachsstock et al., 1994, Berro and Pollard, 2014, Bieling et al., 2016, Gardel et al., 2004). Optimal crosslinking of the actin filaments is thus critical to initiate membrane bending when the force required is the highest. Once membrane bending is initiated, the force required for pulling an invagination likely lowers, and filament nucleation and actin polymerization are therefore sufficient to drive invagination growth in the absence of the crosslinker Sac6. However, when the nucleation of new filaments drops 2 seconds before scission, the mechanical properties of the actin network become important again to provide the force necessary to proceed to the scission of the vesicle: A crosslinked actin network might be able to store and then release the stress accumulated during the first part of the invagination growth, and thus expand and pull the invagination until scission occurs. The release of stress could occur either due to the elasticity of the whole actin network or as a consequence of crosslinker turnover, which could release filament contacts that were induced under higher loads. The enlarged actin network in *bbc1*Δ may accumulate more force than in wild type cells, which could explain the speed-up during the final stage of endocytosis.

## Materials and Methods

### Strain generation

Yeast strains were generated by homologous recombination of the target genes with PCR cassettes encoding for the fluorescent tags (Janke et al., 2004). The C-terminal fluorescent tags were inserted from plasmids pFA6a-EGFP-His3MX6, pFA6a-mCherry-KanMX4 and pYM12-PKS134, which was used for monomeric GFP (myEGFP).

#### Strains used for correlative light and electron microscopy

~~~
*sla2Δ*, Sla1-EGFP, Abp1-mCherry (Skruzny et al., 2012)(MKY1195):
~~~

~~~
MATa, his3Δ200, leu2-3,112, ura3-52, lys2-801, sla2:natNT2, SLA1-EGFP::HIS3MX6, ABP1-mCherry::kanMX4
~~~

~~~
*ent1ΔACB-GFP*, *sla2ΔTHATCH*, Abp1-mCherry (Skruzny et al., 2012)(MKY1846):
~~~

~~~
MATa/α, his3Δ200, ura3-52, lys2-801, ent1ΔACB(Δ amino acids 294–450)-EGFP::HIS3MX6, sla2ΔTHATCH::natNT2, ABP1-mCherry::kanMX4
~~~

~~~
*sac6Δ*, Sla1-EGFP, Abp1-mCherry (Kaksonen et al., 2005)(MKY2869):
~~~

~~~
MATa, his3Δ200, leu2-3,112, ura3-52, lys2-801, sac6:natNT2, SLA1-EGFP::HIS3MX6, ABP1-mCherry::kanMX4
~~~

~~~
*bbc1Δ*, Sla1-EGFP, Abp1-mCherry (Kaksonen et al., 2005)(MKY1805):
~~~

~~~
MATa, his3Δ200, leu2-3,112, ura3-52, lys2-801, bbc1::cgLEU2, SLA1-EGFP::HIS3MX6, ABP1-mCherry::kanMX4
~~~

~~~
*bbc1Δ*, Rvs167-EGFP, Abp1-mCherry (Kaksonen et al., 2005)(MKY1803):
~~~

~~~
MATa, his3Δ200, leu2-3,112, ura3-52, lys2-801, bbc1::cgLEU2, RVS167-EGFP::HIS3MX6, ABP1-mCherry::kanMX4
~~~

#### Strains used for fluorescence live microscopy

~~~
*bbc1∆,* Sla1-EGFP (Kaksonen et al., 2005)(MKY3412):
~~~

~~~
MATα , his3200, leu2-3,112, ura3-52, lys2-801, Sla1-EGFP::HIS3MX6, bbc1delta::natNT2
~~~

~~~
*bbc1∆,* Abp1-EGFP (MKY3247):
~~~

~~~
MATa, his3200, leu2-3,112, ura3-52, lys2-801, Abp1-EGFP::HIS3MX6, bbc1delta::natNT2
~~~

~~~
*bbc1∆,* Rvs167-EGFP (Kaksonen et al., 2005)(MKY3416):
~~~

~~~
MATα , his3200, leu2-3,112, ura3-52, lys2-801, Rvs167-EGFP::HIS3MX6, bbc1delta::natNT2
~~~

~~~
*bbc1∆*, Arc18-myEGFP (MKY3016):
~~~

~~~
MATa, his3200, leu2-3,112, ura3-52, lys2-801, Arc18-myEGFP::NAT, bbc1∆::natNT2
~~~

~~~
*bbc1∆*, Las17-EGFP (MKY3418):
~~~

~~~
MATa, his3200, leu2-3,112, ura3-52, lys2-801, Las17-EGFP::HISMX6, bbc1∆::natNT2
~~~

~~~
*bbc1∆*, Sla1-EGFP, Abp1-mCherry (Kaksonen et al., 2005)(MKY2059):
~~~

~~~
MATa, his3200, leu2-3,112, ura3-52, lys2-801, Sla1-EGFP::HIS3MX6, Abp1-mCherry::KANMX4, bbc1delta::cgLEU2
~~~

~~~
*bbc1∆*, Abp1-mCherry, Rvs167-EGFP (MKY2061):
~~~

~~~
MATa, his3200, leu2-3,112, ura3-52, lys2-801, Rvs167-EGFP::HIS3MX6, Abp1-mCherry, bbc1delta::cgLEU2
~~~

~~~
*bbc1∆*, Arc18-myEGFP, Abp1-mCherry (MKY3015):
~~~

~~~
MATα, his3200, leu2-3,112, ura3-52, lys2-801, Arc18-myEGFP::NAT, Abp1-mCherry::KANMX bbc1∆::natNT2
~~~

~~~
*bbc1∆*, Las17-EGFP, Abp1-mCherry (Kaksonen et al., 2005)(MKY2103):
~~~

~~~
MATα, his3200, leu2-3,112, ura3-52, lys2-801, Las17-EGFP::HIS3MX6, Abp1-mCherry::KANMX, bbc1delta::cgLEU2
~~~

### Live cell imaging

Yeast strains were imaged as described before in (Picco et al., 2015) and (Picco and Kaksonen, 2017). In short, yeast cells were grown overnight at 25ºC in SC-Trp. They were diluted in the morning and grown to log phase at 25ºC. Cells were adhered to 25 mm coverslips (Menzel-Gläser #1), which were coated with Concanavalin A (100 μg/ml). Adhered cells were imaged in SC-Trp medium with an Olympus IX81 microscope and an Olympus 100 x/1.45 NA TIRF objective. Cells were excited for 80-250 ms per exposure with a 488 nm laser or simultaneously with 488 nm and 561 nm lasers. For GFP only images the emission light was filtered using the GFP-3035C-OMF single-band filter set (Semrock, Rochester, NY). For GFP and mCherry two-color simultaneous imaging, the excitation light was reflected with the OBS-U-M2TIR 488/561 (Semrock, Rochester, NY) dichroic mirror and the emission light was split and filtered with the DUAL-view beam splitter (Optical Insights, LLC, Tucson, AZ).

### Image analysis and trajectory alignment

Images were background subtracted and corrected for photobleaching. Before tracking the endocytic events, the cytoplasmic fluorescent signal was removed as described in (Picco et al., 2015) and (Picco and Kaksonen, 2017). Briefly, a median filter of radius 6 pixels was used to estimate the cytoplasmic fluorescent signal, which was then subtracted from the image itself. The endocytic events were tracked using the Particle Tracker plugin in Fiji/ImageJ (Sbalzarini and Koumoutsakos, 2005). All trajectory positions were corrected for chromatic aberrations by applying a warping transformation that was computed from TetraSpeck bead images acquired with the same imaging conditions (Picco et al., 2015). After that, the trajectories of each protein, with the exception of Las17-GFP, were aligned in space and in time and averaged using the trajalign module distribution available in <u>http://apicco.github.io/trajectory_alignment/</u> (Picco and Kaksonen, 2017). Sla1-GFP average trajectories in Figure 2B and C were then aligned in time manually to compare their invagination dynamics. To align the average trajectories of each protein to the other proteins in the same yeast strain, we imaged simultaneously each protein of interest, tagged with EGFP, and Abp1, tagged with mCherry, in both wild-type and *bbc1∆* strains. We then tracked the dynamics and the appearance of the proteins of interest with respect to Abp1 and we used these trajectories to register the position and appearance of each average trajectory with respect to Abp1 (Picco et al., 2015, Picco and Kaksonen, 2017). As Las17-GFP does not move (Picco et al., 2015), Las17 trajectories were tracked simultaneously with Abp1-mCherry trajectories. They were then aligned and averaged using the software in (Picco et al., 2015), which used the spatial and temporal alignment that aligned the Abp1-mCherry trajectories together to compute the transformations that aligned Las17-GFP trajectories. The average trajectories in the different strains were then plotted together in Figure 4 and 6 by using the peak of the Rvs167 number of molecules in both the wild type and in *bbc1∆* to mark the time 0. In Figure 4, the positions along the Y-axis are the positions of the average trajectories in respect to Abp1 shifted so that the average centroid positions of Sla1 trajectories before the invagination dynamics starts are centred on 0.

### Quantification of the number of proteins

The average protein amounts were estimated by comparing the fluorescence intensities of the endocytic spots of the proteins of interest with the fluorescence intensity of Nuf2-GFP spots in cells imaged during their anaphase-telophase (Picco et al., 2015, Joglekar et al., 2006). The average amount of Arc18-myGFP was estimated by comparing to Nuf2-GFP spot intensities and by applying a correction constant of 0.68 ± 0.14 (Picco et al., 2015). The average protein amounts were then used to calibrate the fluorescence intensity curves by scaling the average fluorescence intensity value (Picco et al., 2015).

### Correlative light and electron microscopy

Correlative light and electron microscopy was carried out as described before (Kukulski et al., 2011, Kukulski et al., 2012b). In short, yeast cultures were grown at 25°C in SC-Trp medium to exponential phase. Cells were pelleted using vacuum filtration and high pressure frozen using a Bal-tec HPM010 (McDonald, 2007). Freeze substitution, Lowicryl HM20 embedding and sectioning were done according to the protocol in (Kukulski et al., 2012a). As fiducial markers for the correlation procedure, we used either blue 20 nm FluoSpheres (Molecular Probes) (Kukulski et al., 2012a) or 50 nm TetraSpeck beads (Life Technologies) (Suresh et al., 2015). Fluorescence microscopy of the resin sections on electron microscopy grids was performed immediately after sectioning, as detailed in (Suresh et al., 2015, Kukulski et al., 2012b). Low magnification (2.53 nm pixel size) and high magnification (1.18 nm pixel size) electron tomographic tilt series were acquired on a FEI Tecnai F30 microscope, set up as described in (Kukulski et al., 2012a), using Serial EM (Mastronarde, 2005) and reconstructed with IMOD (Kremer et al., 1996). Centroids of GFP and mCherry signals were correlated to virtual slices of the low magnification electron tomograms based on the fluorescent fiducial makers and in a second step to virtual slices of the high magnification tomograms based on gold fiducial markers using MATLAB-based correlation procedures described in (Kukulski et al., 2011).

### Quantification of membrane shape parameters and ribosome exclusion zones

To quantify shapes of invaginations and vesicles as well as ribosome exclusion zones, we used precisely the same procedures as for the wild type data set (Kukulski et al., 2012a). This allowed us to directly compare the parameters measured in the mutant cells with the wild type data. In short, the Amira EM package (Pruggnaller et al., 2008) was used to click points along the cytoplasmic leaflet of plasma membrane invaginations, in a selected virtual slice of the electron tomogram which was positioned to contain the long axis of the invagination. The groups of points were transferred into a coordinate system such that the x-axis corresponded to the plasma membrane. A local second-degree polynomial was fitted through the points using MATLAB. This fit was used to measure invagination length and radius of the invagination tip, as well as appearance and position of a neck. The Amira EM package was also used to click points along the cytoplasmic leaflet of the vesicle membrane. This was done in the set of virtual slices that comprised the whole vesicle, allowing us to fit an ellipsoid through the set of points. From the major axes of the ellipsoid, we calculated the vesicle surface area. In tomograms that contained vesicles, we also clicked points on the plasma membrane throughout the tomographic volume. We then fitted a sphere through these points and determined the shortest distance of the vesicle centre to the plasma membrane. To quantify the sizes of ribosome exclusion zones, we first generated an average of 10 consecutive virtual slices of which the central slice contained the invagination or the vesicle center. We overlaid this average with a hexagonal mesh of 50 nm spacing and determined which hexagons did not contain ribosomes, as well as which hexagons contained the plasma membrane. These data provided us with estimates for the diameter of the exclusion zone (parallel to the plasma membrane) as well as the height of the exclusion zone. To calculate the volume of the exclusion zones of invaginations, we assumed the shape of a half-ellipsoid with two equal short axes.

### Statistics

For statistical analysis of the membrane shapes and exclusion zones, we used GraphPad Prism or R (<u>https://cran.r-project.org/</u>). All mutant data were compared to wild type data (Kukulski et al., 2012a). For the data that are normally distributed, we used an unpaired two-tailed Welch’s test. To compare the exclusion zone volumes, the vesicle surfaces and their distance from the plasma membrane, we used a Mann-Whitney unpaired nonparametric test. All measurements from CLEM data shown in the figures are listed in Supplemental File 4. Numbers (n) referring to CLEM data set sizes are technical replicates.

### Endocytic sites with multiple invaginations or vesicles

As described for wild type cells, in the correlative microscopy data set of mutant strains described here we found endocytic sites that contained multiple endocytic events within the same exclusion zone (Kukulski et al., 2012a). In the *sac6*Δ data set, one Sla1-GFP Abp1mCherry spots contained two invaginations. These were included in all analysis.

In the *bbc1*Δ data set, of the 26 Rvs167-GFP/Abp1-mCherry spots, 6 contained multiple endocytic events. One site contained an invagination and a vesicle, one contained two invaginations, two contained two vesicles and two contained three vesicles each. Of 25 Sla1GFP/Abp1-mCherry spots containing either invaginations or vesicles, four contained multiple events. One contained a vesicle and an invagination, two contained two invaginations and one contained two vesicles. Of the 14 Abp1-mCherry spots in absence of Sla1-GFP, two contained multiple events. One contained an invagination and a vesicle, and one contained three vesicles. We included these events into all analysis of membrane shape parameters. However, we did not use ribosome exclusion zones that contained multiple events for any quantification of exclusion zone sizes.

## Acknowledgements

This work was supported by the EMBL electron microscopy core facility. A.P. acknowledges an EIPOD fellowship. Work in J.A.G.B’s lab was supported by the Chica und Heinz Schaller Stiftung, the EMBL and the Medical Research Council (MC_UP_1201/16). Work in M.K.’s lab was supported by the Swiss National Science Foundation (31003A_163267). W.K. acknowledges an EIPOD fellowship and postdoctoral fellowships from the Swiss National Science Foundation, as well as support by the Medical Research Council (MC_UP_1201/8).

---

**Supplemental Figure S1:**
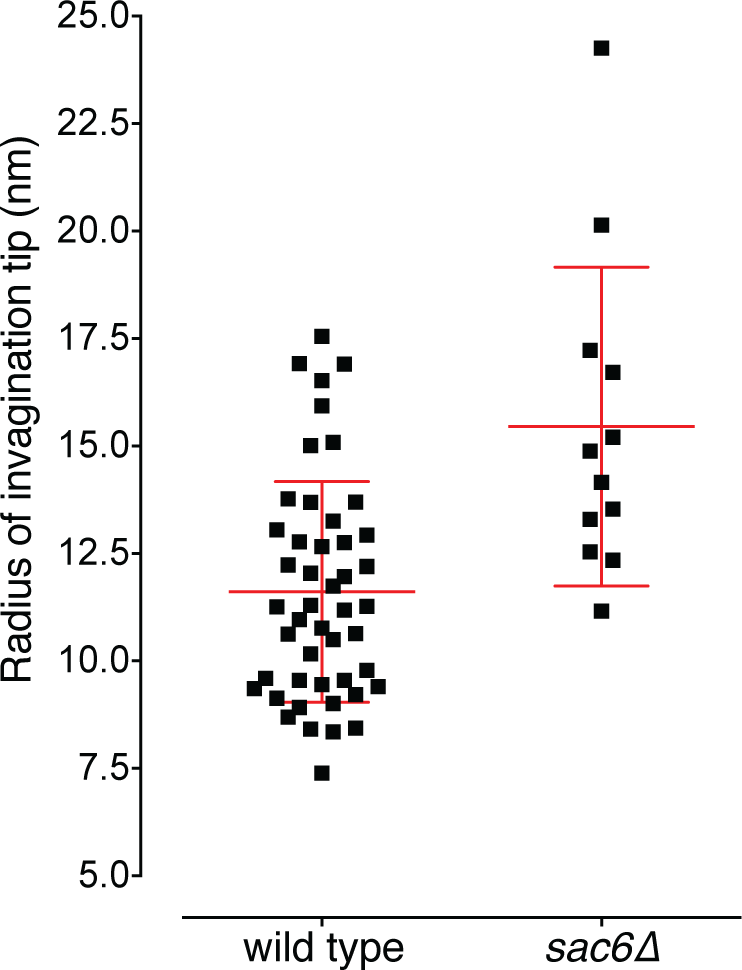
The radii of the tips of the invaginations that were longer than 40 nm, in wild type (Kukulski et al., 2012a) and *sac6Δ* cells. The red lines indicate the mean and the standard deviation.

**Supplemental Figure S2:**
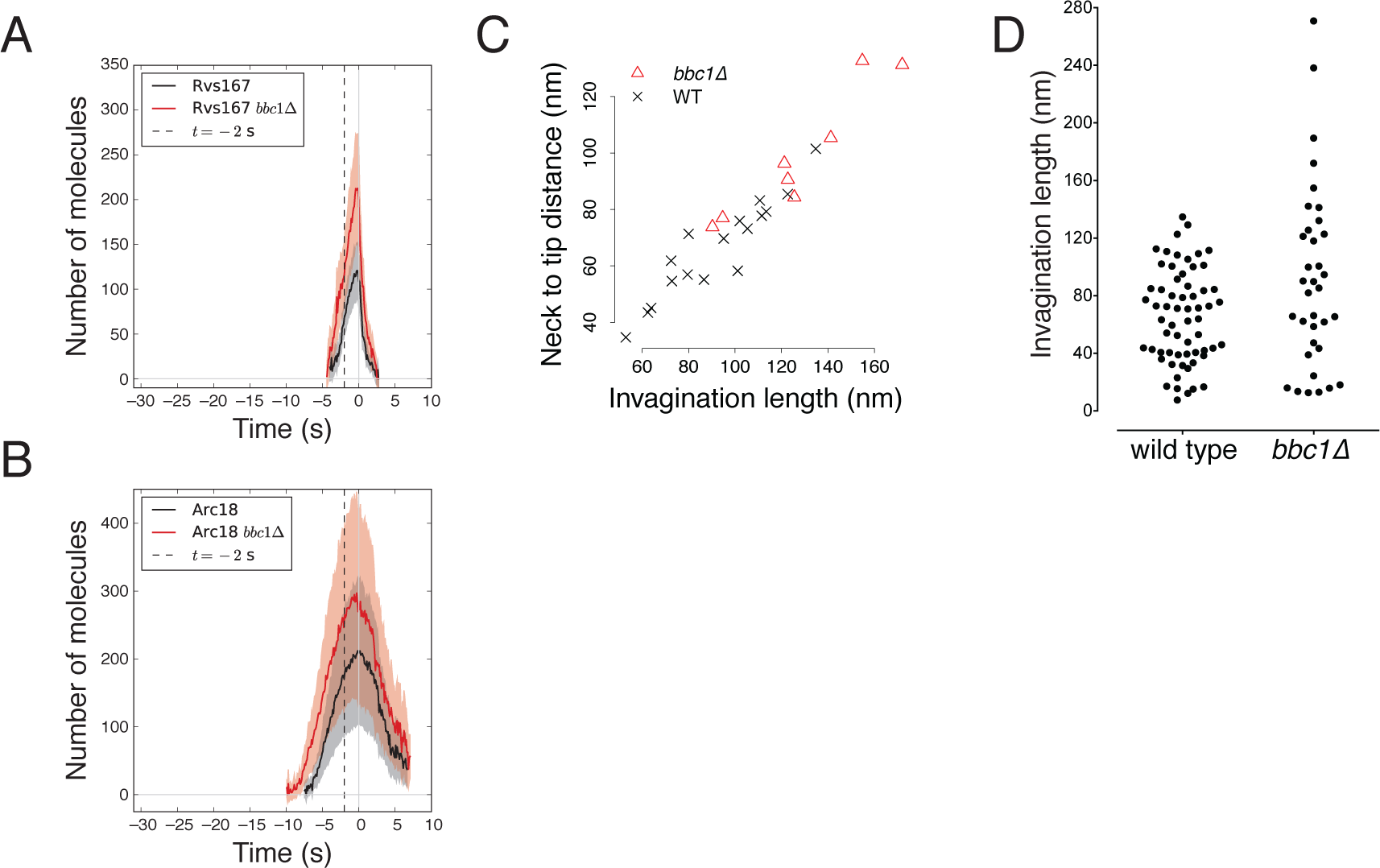
(**A**) The number of Rvs167-GFP molecules that accumulate over time in *bbc1Δ* (red) compared to wild type (black, (Picco et al., 2015)) cells. (**B**) The number of molecules of Arc18-GFP in *bbc1Δ* (red) and in wild type (black, (Picco et al., 2015)) cells. In A and B, the trajectories were independently aligned in space and in time to Abp1 (Picco and Kaksonen, 2017). The time point 0 marks the scission event (Kukulski et al., 2012a, Picco et al., 2015) (**C**) The position of invagination necks in *bbc1Δ* (red triangles) and in wild type (black crosses (Kukulski et al., 2012a)) cells. (**D**) The invagination lengths in *bbc1Δ* compared to the invagination lengths in wild type cells (Kukulski et al., 2012a).

**Supplemental File S3:** Table containing the data points of the average trajectories shown in Figures 2B and C, Figure 4A, B, C and D, Figure 6, as well as Supplemental Figure S2A and B.

**Supplemental File S4:** Table containing the data points of measurements of membrane shapes and exclusion zones, shown in Figures 3C, Figure 5C, D, E and F, Supplemental Figure S1, as well as Supplemental Figure S2C and D.

